# Assessing 42 inflammatory markers in 321 control subjects and 887 major depressive disorder cases: BMI and other confounders and overall predictive ability for current depression

**DOI:** 10.1101/327239

**Authors:** Timothy R. Powell, Helena Gaspar, Raymond Chung, Aoife Keohane, Cerisse Gunasinghe, Rudolf Uher, Katherine J. Aitchison, Daniel Souery, Ole Mors, Wolfgang Maier, Astrid Zobel, Marcella Rietschel, Neven Henigsberg, Mojca Zvezdana Dernovšek, Joanna Hauser, Souci Frissa, Laura Goodwin, Matthew Hotopf, Stephani L Hatch, David A. Collier, Hong Wang, Hong Wang

**Author notes:** Corresponding Author: Dr Gerome Breen, Social, Genetic and Developmental Psychiatry Centre, Institute of Psychiatry, Psychology and Neuroscience, King’s College London, PO80, 16 De Crespigny Park, London, SE58AF, UK.

## Abstract

Inflammatory markers such as cytokines represent potential biomarkers for major depressive disorder (MDD). Many, generally small studies have examined the role of single markers and found significant associations. We assessed 42 inflammatory markers, namely cytokines, in the blood of 321 control subjects and 887 MDD cases. We tested whether individual inflammatory marker levels were significantly affected by MDD case/control status, current episode, or current depression severity, co-varying for age, sex, body mass index (BMI), smoking, current antidepressant use, ethnicity, assay batch and study effects. We further used machine learning algorithms to investigate if we could use our data to blindly discriminate MDD patients, or those in a current episode. We found broad and powerful influences of confounding factors on log-protein levels. Notably, IL-6 levels were very strongly influenced by BMI (p = 1.37 × 10^−43^, variance explained = 18%), while Interleukin-16 was the most significant predictor of current depressive episode (p = 0.003, variance explained = 0.9%, q < 0.1). No single inflammatory marker predicted MDD case/control status when a subject was not in a depressed episode, nor did any predict depression severity. Machine learning results revealed that using inflammatory marker data with clinical confounder information significantly increased precision for differentiating MDD patients who were in an episode. To conclude, a wide panel of inflammatory markers alongside clinical information may aid in predicting the onset of symptoms, but no single inflammatory protein is likely to represent a clinically useful biomarker for MDD diagnosis or prognosis. We note that the potential influence of physical health related and population stratification related confounders on inflammatory biomarker studies in psychiatry is considerable.

## Introduction

Inflammatory proteins, such as cytokines, are mediators of the immune system and are key in orchestrating appropriate responses to infection.^1–3^ Pro-inflammatory cytokines promote systemic inflammation and are predominantly released by macrophages and T-cells.^4–5^ Interleukin-6 (IL-6) and tumour necrosis factor (TNF) are key examples of pro-inflammatory cytokines released endogenously to combat infection.^4^ Chemokines are a subset of smaller cytokines (e.g. interleukin-8), which act as chemotactic factors, and help to direct immune cells, such as neutrophils, to the site of infection, where they can aid in eliminating a pathogen.^1^ In contrast, anti-inflammatory cytokines work to reduce inflammation and are important in shutting down the pro-inflammatory state, to assist in wound healing; a key example is interleukin-10 (IL-10) which is primarily synthesized by monocytes.^7–8^ The general inflammatory status of an individual is often characterised clinically by C-reactive protein (CRP), which is an acute phase protein synthesized in the liver, which rises in response to inflammation.^6^

In addition to its important role in combatting infection, a pro-inflammatory profile is reportedly associated with a broad range of disease states including diabetes, obesity and cancer.^9–10^ Studies have also reported increased circulating levels of pro-inflammatory cytokines in psychiatric disorder patients, and it has been proposed that cytokines influence neurotransmitter systems and brain functionality related to psychiatric disease pathology.^11^ Previous studies of major depressive disorder patients (MDD) have shown heightened circulating levels of pro-inflammatory cytokines and acute phase proteins, including IL-6, TNF and CRP, and lower levels of the anti-inflammatory cytokine IL-10.^12–15^ In the cases of IL-6, TNF and IL-10, the body of associative evidence with MDD has been drawn from small case-control studies (mainly n < 50) and meta-analyses, many of which only account for the confounding effects of age and gender.^16–20^

Body mass index (BMI), smoking and ethnicity have been found to influence, in some cases very strongly, the circulating levels of cytokines and represent very plausible confounding factors in MDD case-control studies and within-case studies.^25–27^ Especially considering, MDD patients have an increased tendency to smoke, chronic cases having a higher BMI, and MDD prevalence varies amongst different ethnicities.^21–24^ Furthermore, cytokines and acute phase proteins are heavily influenced by each other’s expression^28^, and few studies have considered whether cytokines in the wider inflammatory pathway may have a more pervasive association with MDD, than just IL-6 and CRP.

The examination of case-control differences in inflammatory marker (e.g. cytokines) expression within well-characterised cohorts could be useful for two reasons. First, confirmed differences in the expression of specific cytokines, may hint toward an immuno-inflammatory pathophysiology of MDD, allowing for the creation of drugs targeting specific components of the immune system in order to treat it.^29^ Secondly, differences in cytokine levels may provide an objective test for an individual’s diagnosis and prognosis. For instance, if heightened cytokine levels precede a severe depressive episode, a cytokine biomarker might allow for treatment to be initiated more rapidly, potentially reducing the risk of suicide.^11^

In this study we investigated the utility of 42 inflammatory proteins, namely cytokines, as biomarkers for the prediction of MDD case-control status, current depressive episode, and episode severity, utilising blood serum from 321 control subjects and 887 MDD cases, making this one of the most extensive studies of its kind to-date. We further studied the effects of age, sex, BMI, current smoking, ethnicity, and current antidepressant medication on circulating levels of cytokines, and we used a machine learning approach to examine if inflammatory markers in conjunction with clinical/confounder information could increase the precision of blind MDD discrimination when patients were, and were not, in a depressed episode.

## Methods

### The Study Sample

Peripheral blood samples utilized here were obtained by venipuncture as part of four separate studies: SELCoH, HDAO (ClinicalTrials.gov Identifier:NCT00035321), LNBI (ClinicalTrials.gov Identifier:NCT00795821) and GENDEP. All serum was stored at −80 °C prior to use. Subject information relating to each study is shown in Table 1 and a description of each study is given below.

### SELCoH

The South East London Community Health Study (SELCoH) is a population study in London, UK, investigating community health. Within this sample, there were samples collected from 27 families (total n=75) as well as 420 non-related individuals. So far, participants have undergone extensive and repeated phenotypic assessment as part of three separate phases. In the third phase, biological specimens were collected from a subset of participants, which included blood for serum separation.^30–33^ MDD case/control status was characterised using the Clinical Interview Schedule-Revised (CIS-R), which can be used to generate ICD-10 diagnoses.^34^ A participant was screened positive for an MDD diagnosis if the CIS-R identified a moderate-to-severe depressive episode in any one of the three interviews. A case was screened positive for a current depressive episode if they were in a moderate-to-severe depressive episode in the third phase (i.e. when blood was collected). 321 control subjects within SELCoH were identified as those with no depression symptoms during any of the three interviews, with no previous diagnosis of a depressive disorder (based on self-report). We further identified a subset of 257 ‘super controls’ who showed no psychiatric symptoms at all (i.e. outside of depression) in any of the three interviews, and no previous history of depression; this subset of controls was used for secondary analyses in a more stringent case-control comparison.

### HDAO

The HDAO study was a clinical trial carried out in the United States by Eli Lilly testing the differential effects of fluoxetine, olanzapine, and fluoxetine + olanzapine combinations on therapeutic response in MDD patients.^35^ Eligibility criteria included: meeting DSM-IV criteria for recurrent MDD without psychotic features^36^, a current depressive episode, and previous failure to achieve response to an antidepressant other than a selective serotonin reuptake inhibitor. Blood serum was collected from patients at baseline, which we utilise in this study.

### LNBI

The LNBI study was a clinical trial carried out in the United States by Eli Lilly testing the effect of the antidepressant edivoxetine (LY2216684) relative to placebo on major depression symptoms.^37^ Eligibility criteria included DSM-IV-TR criteria for MDD without psychotic features^36^, and a current depressive episode as assessed using the Clinical Global Impressions Scale (CGIS).^38^ Depression severity was captured using the Montgomery Asberg Depression Rating Scale (MADRS).^39^

### GENDEP

The Genome-based Therapeutics Drugs for Depression (GENDEP) was a 12-week partially randomized open label pharmacogenetic study in European MDD patients, comparing the effects of escitalopram and nortriptyline on symptom improvements.^28–29,40–44^ The study consisted of treatment-seeking adults with MDD symptoms of at least moderate severity according to ICD-10 or DSM-IV criteria.^36,45^ Blood serum was collected from patients at baseline, which we utilise in this study. Depression severity was characterized using three rating scales, including the MADRS.^39^

**Table 1:** A summary of characteristics within the four subject cohorts included in this study.

| Study | n | Case-Control Status | Age | Sex (% male) | BMI | Ethnicity | Current Episode (n) | Current Antidepressant (n) | Current Smoker (n) |
| --- | --- | --- | --- | --- | --- | --- | --- | --- | --- |
| SELCoh | 321 | Controls | 49.91 (16.19) | 46.4 | 26.96 (5.23) | Black (20.9%) | 0 | 0 | 52 |
|  |  |  |  |  |  | Other (3.7%) |  |  |  |
|  |  |  |  |  |  | White (75.4%) |  |  |  |
| SELCoh | 69 | Cases | 45.62 (14.70) | 34.8 | 28.77 (6.50) | Black (18.8%) | 38 | 28 | 28 |
|  |  |  |  |  |  | Other (5.8%) |  |  |  |
|  |  |  |  |  |  | White (75.4%) |  |  |  |
| HDAO | 488 | Cases | 43.77 (10.20) | 34.6 | 30.17 (6.98) | Black (4.3%) | 488 | 40 | - |
|  |  |  |  |  |  | Other (14.3%) |  |  |  |
|  |  |  |  |  |  | White (81.4%) |  |  |  |
| LNBI | 128 | Cases | 44.61 (11.24) | 54.7 | - | Black (21.9%) | 128 | 1 | 87 |
|  |  |  |  |  |  | Other (0%) |  |  |  |
|  |  |  |  |  |  | White (78.1%) |  |  |  |
| GENDEP | 202 | Cases | 41.50 (12.45) | 42.6 | 25.22 (5.06) | Black (0%) | 202 | 0 | 48 |
|  |  |  |  |  |  | Other (0%) |  |  |  |
|  |  |  |  |  |  | White (100%) |  |  |  |

### Inflammatory Marker Quantification

Upon use, serum was thawed at room temperature and 42 inflammatory markers were quantified simultaneously using multiplex ELISA-based technology provided by the Meso Scale Discovery V-PLEX Plus Human Biomarker 40-Plex kit, and a customised human duplex kit assaying brain-derived neurotrophic factor (BDNF) and interferon-alpha (IFN-α). Note however, that interleukin-8 (IL-8) is repeated twice on the 40-plex array (IL-8 and IL-8(HA)) alongside two different standard curves, allowing for a very wide range of IL-8 levels to be detected. We only utilized data from IL-8 (not IL-8(HA)) as our samples were detectable specifically within the range of this standard curve (0.0700 - 498 pg/mL). The 42 capture antibodies are etched to the bottom of five 96-well plates, each capturing between 2 and 10 inflammatory markers. Seven-point standard curves were run in duplicate on each plate in order to calculate absolute pg/mL values for the 80 samples assayed per plate, and a no-template control was used to correct for background fluorescence. Cases and controls were randomised across batches, and plates were scanned on the Mesoscale Scale Discovery MESO Quickplex SQ 120 reader at the MRC SGDP Centre, Institute of Psychiatry, Psychology and Neuroscience, King’s College London. Pilot studies revealed very high intra-plate (r > 0.99) and interplate (r > 0.97) correlations, suggesting single measurements were acceptably reliable using this methodology. Furthermore, known quantities within the standard curves used on each plate, correlated very highly with quantities predicted by fluorescence intensity (r > 0.99). We additionally validated our methodology by comparing results obtained using an independent method (single ELISA for C-reactive protein) in a independent laboratory in the same sample set ^43^, and our results showed a high positive correlation (r > 0.85).

### Statistical Analysis

#### (i) Data Processing

Standard curves were used to determine absolute quantities (pg/mL) of each inflammatory marker. Absolute quantities (pg/mL) were then log-transformed to allow for parametric analyses. We excluded inflammatory markers where greater than 20% of data was missing, as it indicates that these proteins may not be useful as biomarkers. Subsequently, data points were removed if they exceeded +/− 2 standard deviations from the mean.

#### (ii) Identification of marker-specific confounders

For each log-protein level, we performed a general linear model to test for the effects of age, sex, BMI, ethnicity, family ID (related to the presence of families in SELCoH) and smoking, alongside batch and study. In a systematic, step-wise fashion we dropped any non-significant covariates (p > 0.05), until we established a list of significant confounders to include in our downstream analyses.

#### (iii) Case-control & Current Episode analysis

For the case/control comparison, we performed a general linear regression with log-protein level as the dependent variable, major depression case/control status, current depressive episode, and current antidepressant medication as the independent variables, alongside the following covariates: batch/plate effects, study effects, and any other significant confounders identified from (ii). We applied the false discovery rate for multiple testing correction and a q < 0.1 threshold to determine true associations.^46^ As a secondary analysis, we also performed the same analysis in a subset of super controls (n=257), who were free from all psychiatric symptoms.

#### (iv) Depression Severity Analyses

To test the effect of current depression severity on inflammatory marker levels we performed a within-cases analysis utilising the LNBI and GENDEP cohorts, and MADRS scores. We performed a general linear regression with log-protein level as the dependent variable, MADRS scores as the independent variable, alongside batch/plate effects, study effects, and any other significant confounders. We applied the false discovery rate of multiple testing correction and a q < 0.1 threshold to determine true associations.

#### (v) Machine Learning

Finally, to investigate the collective predictive capabilities of inflammatory marker levels on (i) MDD case/control and (ii) current depressive episode, we used machine learning. Initially, we selected only individuals who were medication-free. Only samples with defined values for age, smoking status, BMI, ethnicity and gender were kept. Missing values were imputed using a k nearest neighbours approach with number of neighbours k = 3. For the remaining inflammatory markers, we regressed out the effects of nuisance factors (batch, study) by taking the standardized residuals. For each phenotype (disease status, current episode), we built three sets of variables for the 463 instances: inflammatory markers, “confounders” (age, smoking status, BMI, ethnicity, gender), and inflammatory markers + confounders.

Each set of variables were separated into training (n=324) and validation sets (n=139). Each training set was used to build several machine learning models evaluated in a 10-fold cross-validation procedure repeated 10 times. The methods included a Naïve Bayes classifier with kernel estimator, Random Forests with 500 or 1000 trees, both implemented in WEKA. The classifiers are cost-sensitive: the training instances were reweighted to account for class imbalance. We selected models with the best precision (Random Forests with 1000 trees, for all sets of variables). This “best” model was then validated on the blinded external validation set. The attribute evaluation method was also performed on the training set to rank the contribution of each variable in contributing to case/control or current episode discrimination. This was achieved using “ReliefF” implemented in WEKA; instances are sampled randomly and the value of the attribute of the nearest instance of the same and different class is considered.^47^

## Results

### (i) Inflammatory markers adequately detected in serum using our methodology

32 inflammatory proteins passed our quality control criteria. 10 inflammatory markers showed greater than 20% missingness across our samples and were removed from downstream analysis (IL4, MIP-1A, IFN-a, GM-CSF, IL-1A, IL-13, IL-1B, IL-2, IL-5, IL-8(HA)), Figure 1 shows all inflammatory markers expressed in >80% of our sample. Inter-plate co-efficient of variation (CV) was calculated, which revealed low levels of inter-plate variability for every marker assessed (CV < 2). See S1 in Supplementary information for details on how the inflammatory markers correlate with one another.

**Figure 1:**
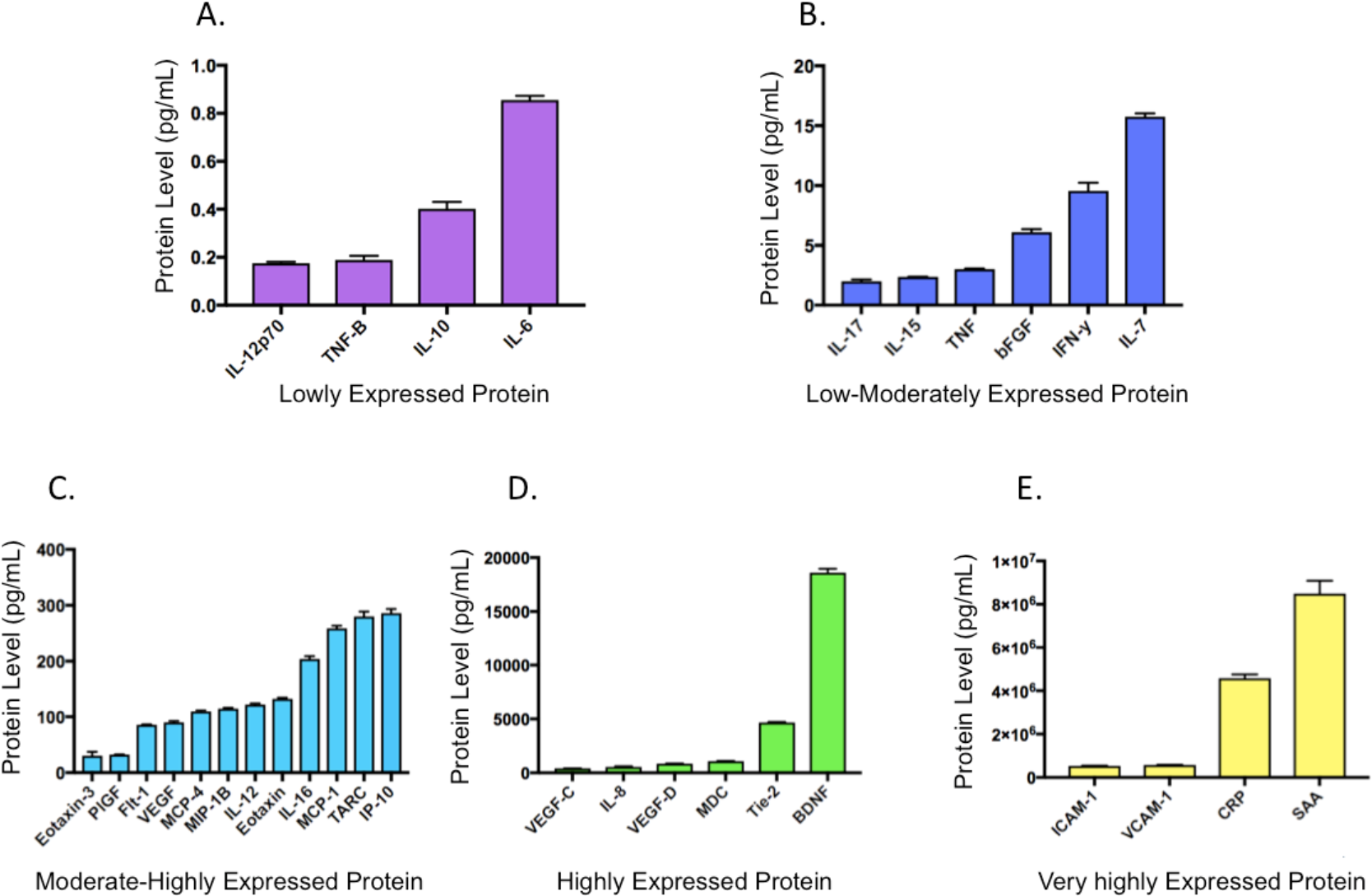
32 inflammatory markers were adequately expressed in our patient sample, and therefore could potentially be used as biomarkers. Figure 1: A summary of inflammatory markers lowly expressed in our sample (A; < 1 pg/mL), low-moderately expressed in our sample (B; 1-25 pg/mL), moderate-highly expressed in our sample (C; 30-300 pg/mL), highly expressed in our sample (D; 400–20,000 pg/mL) and very highly expressed in our sample (E; 500,000–1,000,000,000 pg/mL). Bars represent the mean and error bars represent the standard error of the mean. Inflammatory markers in 1A include: Interleukin 12 heterodimer (IL-12p70); Tumour necrosis factor-beta (TNF-B), interleukin-10 (IL-10), interleukin-6 (IL-6). Inflammatory markers in 1B include: interleukin-17 (IL-17), interleukin-15 (IL-15), tumour necrosis factor (TNF), Basic fibroblast growth factor (bFGF), Interferon gamma (IFN-γ) and interleukin-7 (IL-7). Inflammatory markers in 1C include: Chemokine (C-C motif) ligand 26 (Eotaxin-3), Phosphatidylinositol-glycan biosynthesis class F protein (PIGF), Vascular endothelial growth factor receptor 1 (Flt-1), Vascular endothelial growth factor (VEGF), Monocyte chemoattractant protein 4 (MCP-4), Macrophage Inflammatory Protein β (MIP-1B), interleukin-12 (IL-12), eotaxin-1 (eotaxin), interleukin-16 (IL-16), Monocyte chemoattractant protein 1 (MCP-1), Chemokine (C-C motif) ligand 17 (TARC) and Interferon gamma-induced protein 10 (IP-10). Inflammatory markers in 1D include: Vascular endothelial growth factor C (VEGF-C), interleukin-8 (IL-8), Vascular endothelial growth factor D (VEGF-D), Macrophage-derived chemokine (MDC), Tyrosine kinases with Ig and EGF homology domains-2 (Tie-2), brain-derived neurotrophic factor (BDNF). Inflammatory markers in 1E include: Intercellular Adhesion Molecule 1 (ICAM-1), Vascular cell adhesion molecule 1 (VCAM-1), C-reactive protein (CRP) and Serum amyloid A (SAA).

### (ii) Identification of inflammatory marker-specific confounders

Our results revealed extensive influences of confounding factors on the circulating levels of specific inflammatory markers (with the exception of family ID which did not significantly influence the expression of any inflammatory marker, p > 0.05), Figure 2.

**Figure 2:**
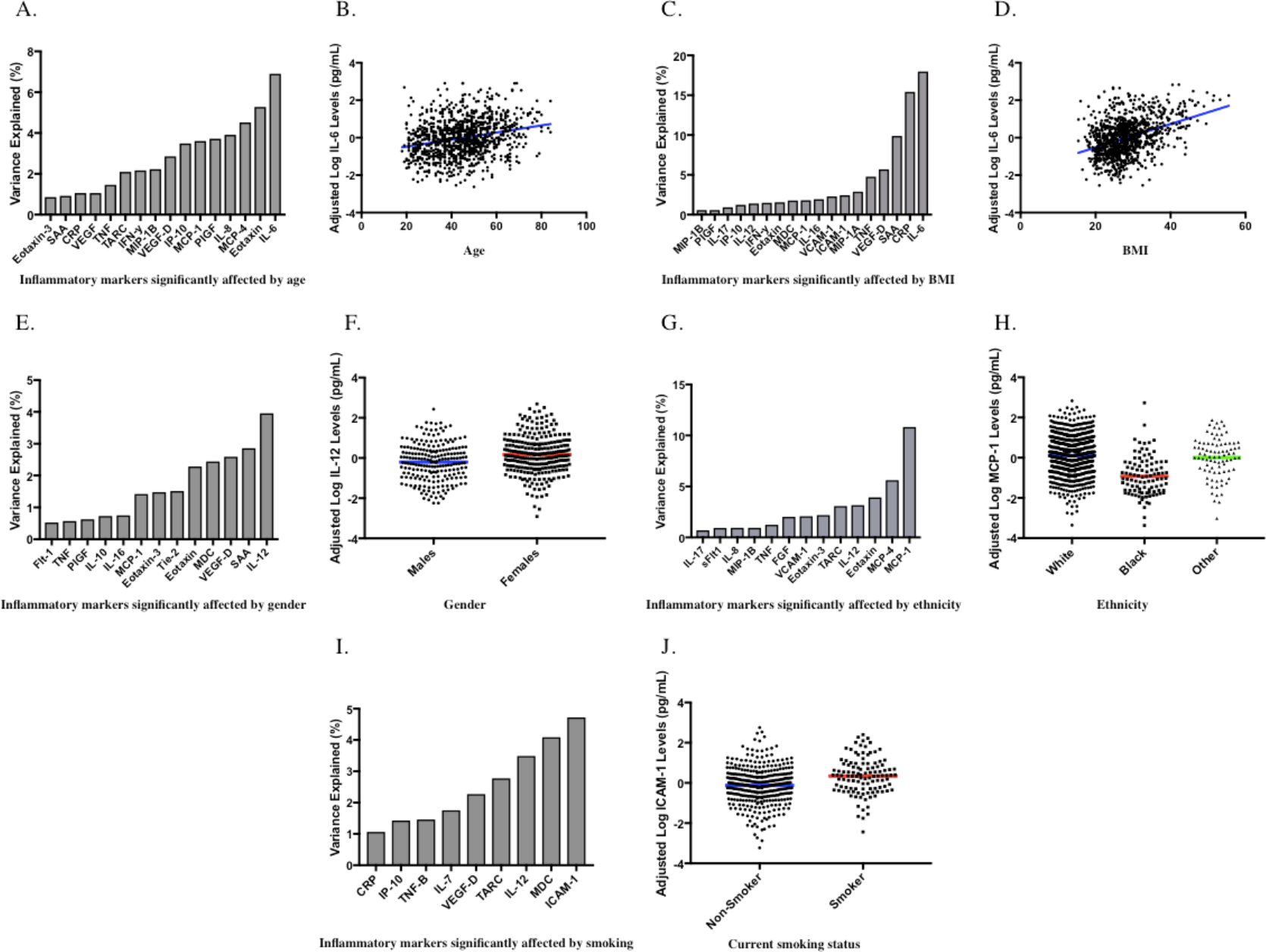
Confounding factors have broad and powerful influences on inflammatory marker expression. Figure 2A: A chart displaying inflammatory markers significantly affected by age on the x-axis, with the percentage of variance explained by age on the y-axis. Figure 2B: A scatterplot showing the relationship between age (x-axis) and adjusted log IL-6 (pg/mL) levels (y-axis), with a line of best fit shown in blue. Results from our linear regression analysis revealed age very strongly predicted IL-6 levels (F(1, 968) = 71.760, p = 8.947 × 10^−17^, variance explained = 6.902%). Log IL-6 data points were adjusted for the effects of batch, study, MDD case/control status, current episode, current antidepressant use and BMI. Figure 2C: A chart displaying inflammatory markers significantly affected by BMI on the x-axis, with the percentage of variance explained by BMI on the y-axis. Figure 2D: A scatterplot showing the relationship between BMI (x-axis) and adjusted log IL-6 (pg/mL) levels (y-axis), with a line of best fit shown in blue. Results from our linear regression analysis revealed BMI very strongly predicted IL-6 levels (F(1, 968) = 212.086, p = 1.371 × 10^−43^, variance explained = 17.972%). Log IL-6 data points were adjusted for the effects of batch, study, MDD case/control status, current episode, current antidepressant use and age. Figure 2E: A chart displaying inflammatory markers significantly affected by gender on the x-axis, with the percentage of variance explained by gender on the y-axis. Figure 2F: A plot showing gender (x-axis) and adjusted log IL-12 pg/mL levels (y-axis), the mean in each group is shown with coloured lines. Results from our linear regression analysis revealed gender significantly predicted IL-12 levels (F(1, 448) = 17.703, p = 3.100 × 10^−5^, variance explained = 3.801%). Log IL-12 data points were adjusted for the effects of batch, study, MDD case/control status, current episode, current antidepressant use, BMI, ethnicity and smoking. Figure 2G: A chart displaying inflammatory markers significantly affected by ethnicity on the x-axis, and the percentage of variance explained by ethnicity on the y-axis. Figure 2H: A plot showing ethnicity (x-axis) and adjusted log MCP-1 pg/mL levels (y-axis), the mean in each group is shown with coloured lines. Results from our linear regression analysis revealed ethnicity significantly predicted MCP-1 levels (F(2, 998) = 59.998, p = 2.559 × 10^−25^, variance explained = 10.830%). Log MCP-1 data points were adjusted for the effects of batch, study, MDD case/control status, current episode, current antidepressant use, gender, age and BMI. Figure 2I: A chart displaying inflammatory markers significantly affected by current smoking status on the x-axis, with the percentage of variance explained by current smoking on the y-axis. Figure 2J: A plot showing smoking status (x-axis) and adjusted log ICAM-1 pg/mL levels (y-axis), the mean in each group is shown with coloured lines. Results from our linear regression analysis revealed current smoking significantly predicted ICAM-1 levels (F(1, 448) = 22.216, p = 3.000 × 10^−6^, variance explained 4.725%). Log ICAM-1 data points were adjusted for the effects of batch, study, MDD case/control status, current episode, current antidepressant use and BMI.

### (iii) Case-control analysis, current episode and antidepressant treatment

The case-control analysis revealed that no inflammatory markers were significantly associated with MDD case-control status, either nominally or after multiple testing correction (q > 0.1), this included the highly cited IL-6 and CRP associations with MDD, Figure 3. There were however four inflammatory markers nominally associated with a current depressive episode (VEGF, FGF, SAA, IL-16); all of which were more highly expressed during a depressed episode. IL-16 was the only inflammatory marker significantly associated with current depression episode after multiple testing correction (F(1, 988) = 8.856, p = 2.992 × 10^−3^, q = 0.096, variance explained = 0.888%), Figure 3, see Supplementary Information S2 for full results. Those currently taking antidepressants had nominally higher circulating levels of CRP (F(1, 456) = 4.106, p = 0.043, variance explained = 0.893%), but no other inflammatory markers were affected by current antidepressant medication. When using our ‘super control’ group relative to MDD cases, our results did not reveal any further associations, see Supplementary Information S3 for full results.

**Figure 3:**
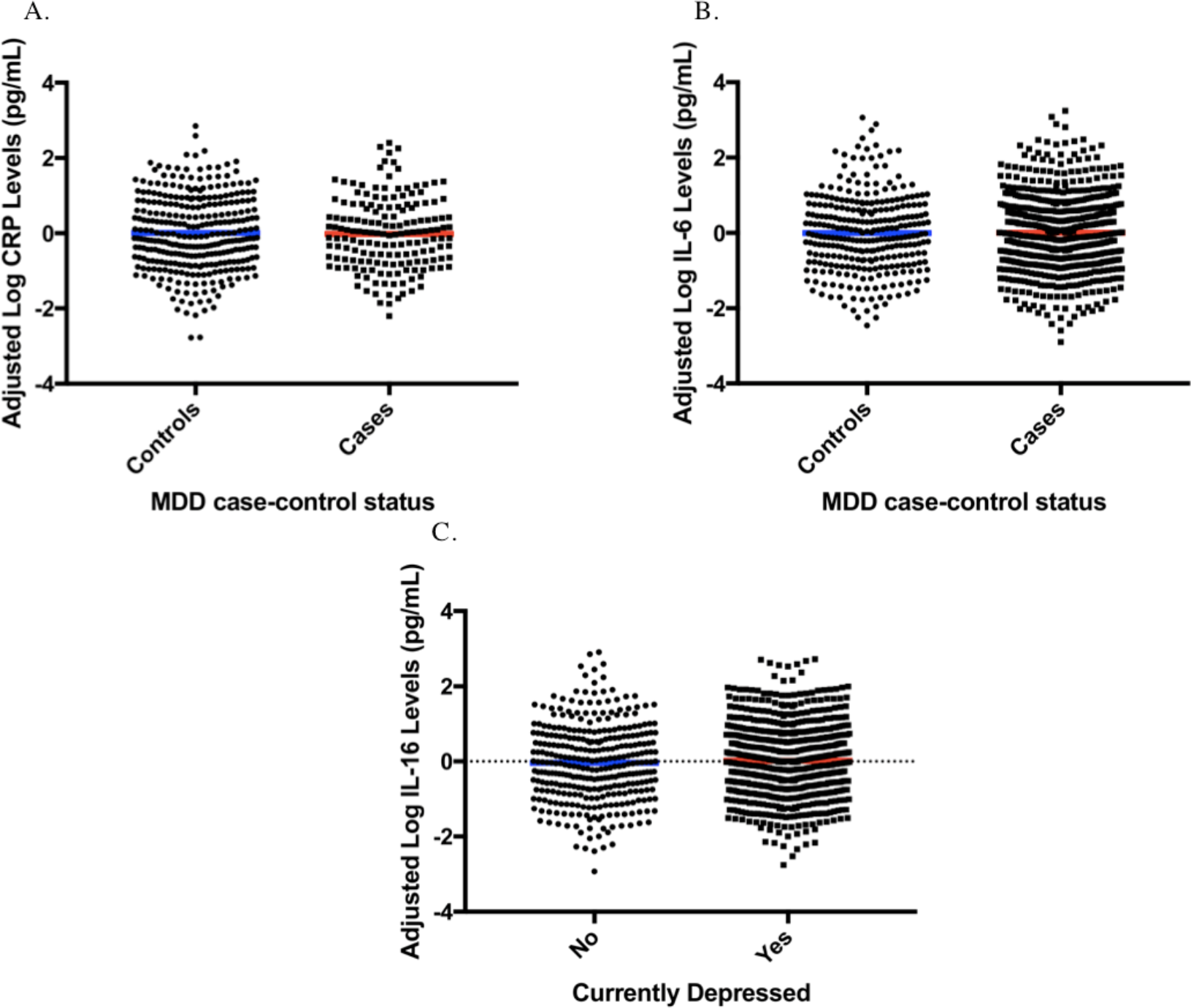
‘Classic’ inflammatory markers do not show an association with MDD, but IL-16 shows a small, yet significant association with current depressed episode. 3A: A plot showing MDD case/control status (x-axis) and adjusted log CRP pg/mL levels (y-axis), the mean in each group is shown with coloured lines. Log CRP data points were adjusted for the effects of batch, study, current episode, current antidepressant use, age, BMI and smoking. Results from our linear regression analysis revealed no differences between groups in the expression of CRP (p>0.05). Figure 3B: A plot showing MDD case/control status (x-axis) and adjusted log IL-6 pg/mL levels (y-axis), the mean in each group is shown with coloured lines. Log IL-6 data points were adjusted for the effects of batch, study, current episode, current antidepressant use, age and BMI. Results from our linear regression analysis revealed no differences between groups in the expression of IL-6 (p>0.05). Figure 3C: A plot showing current depressive episode status (x-axis) and adjusted log IL-16 pg/mL levels (y-axis), the mean in each group is shown with coloured lines. Results from our linear regression analysis revealed current depressive episode nominally predicted IL-16 levels (F(1, 948) = 8.497, p = 0.00364, variance explained 0.9%). Log IL-16 data points were adjusted for the effects of batch, study, MDD case/control status, current antidepressant use, gender and BMI.

### (iv) Depression Severity

The inflammatory marker PIGF was nominally associated with depression severity (F(1, 170) = 4.052, p = 0.046, variance explained = 2.6%). No inflammatory markers were significantly affected by depression severity after multiple testing correction (q > 0.1), see Supplementary Information for full results.

### (v) Machine learning

Models were built to predict current episode or disease status separately. Random Forest (1000 trees) models gave the best results. The cross-validated averaged precision with 95% confidence interval for the best model is given in Figure 4, alongside validation in an external dataset. The best model for disease status as well as current episode is obtained with inflammatory markers + confounders. Current episode is better predicted than disease status. The results indicate that the levels of inflammatory markers provide information on depression not solely due to confounders. The contribution of each variable in aiding the discrimination of cases/controls and currently depressed/not depressed subjects as part of the machine learning process (i.e. ReliefF results) is shown in Figure 5. For Full ReliefF results, see Supplementary information, S4.

**Figure 4:**
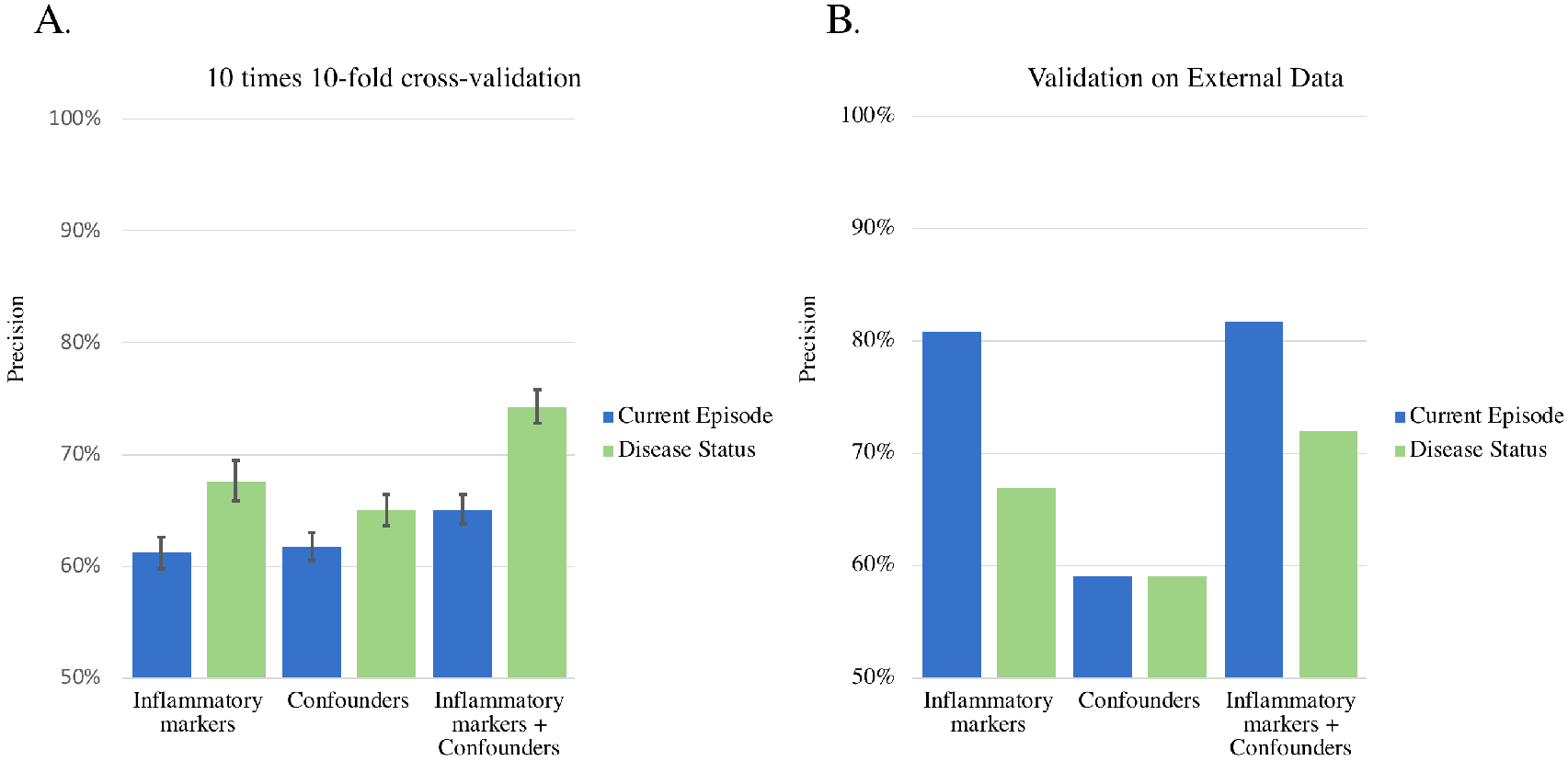
Machine learning results show inflammatory marker information significantly increases the precision of blind MDD diagnoses, especially when a patient is in a current episode. Figure 10: Precision results for the “best” Random Forest model predicting case/control status and current depressed episode using a 10-fold cross-validation repeated 10 times (training set), with 95% confidence interval, (A). Also displayed are the results from an independent sample (validation set; B). In both, the test and replication datasets, the inflammatory markers + confounders group outperformed the confounder group alone, particularly in relation to current depressed episode.

**Figure 5:**
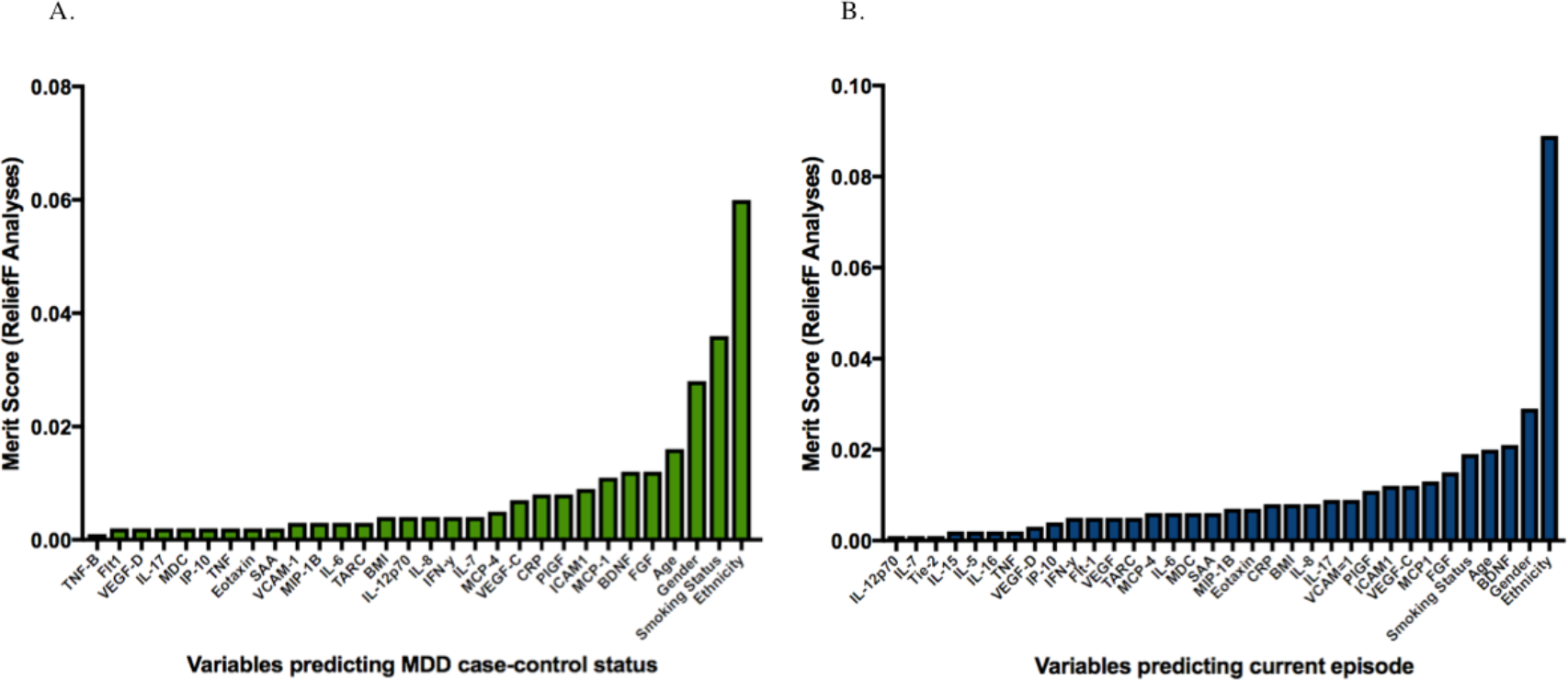
The ranked contribution of variables that increased the precision of MDD case-control discrimination and current episode discrimination as part of the machine learning process. Results from ReliefF, whereby the contribution of each variable was ranked and a ‘merit score’ was generated. We show variables with positive merit scores relating to MDD case-control status (left) and current depressed episode (right).

## Discussion

There has been much research addressing the potential relationship between a heightened pro-inflammatory state and MDD. This study was designed as a well-powered replication of the findings reported by smaller case-control studies, which reported differences in levels of cytokines such as IL-6, amongst cases.^16–20, 48^ Our study represents one of most extensive to-date, investigating differences in the expression of 42 inflammatory markers in relation to MDD case-control status, current episode and depression severity in a large cohort of 1,208 individuals. From our study we can draw four main conclusions.

First, in our study, none of the 42 inflammatory markers investigated, represent biomarkers for lifetime presence of MDD, this included CRP or IL-6, *Figure 3*. Given the size of the study, it appears unlikely that single inflammatory measures could be used prospectively to identify people at risk for developing MDD, who have not yet exhibited symptoms. Second, we found IL-16 levels to be associated with a current depressed state (even after multiple testing correction). If further studies showed that this cytokine increased in expression preceding clinical symptoms, it could be useful in initiating rapid treatment. However, the amount of variance explained by IL-16 is minimal (<1%) and therefore this cytokine is unlikely to be clinically useful. It may, however, represent a component of a larger immuno-inflammatory mechanism related to the precipitation of symptoms which is independent from the cytokines we assayed. Further research is required to understand the influence of IL-16 on depression mechanisms and the brain.

Our first two findings show a sharp contrast with the older and highly cited research performed in smaller samples, demonstrating links between cytokines and MDD^12, 15–17^, but our results do corroborate the findings of more recent studies performed in larger sample sets. For instance, a recent MDD case-control study assaying 18 cytokines (which included IL-6 and TNF) in 236 individuals revealed no significant differences between cases and controls.^50^ Our findings are also reminiscent of the extensive case-control association work performed on functional genetic variants in psychiatric candidate genes (e.g. SERT and COMT), many of which were later disproved as risk factors in far larger, more powerful GWAS.^49^

Our third main finding was that few MDD cytokine case-control studies have been adequately powered to consider the possible confounding effects of environmental factors considerably more frequent amongst MDD patients, such as smoking or higher BMI.^21,22^ We studied the influences of these factors in the current study as well as sex, age, ethnicity, and current antidepressant use. Our study very clearly highlights the extensive effects confounding factors have on the expression of inflammatory markers, *Figure 2*. Most notable are the influences of BMI on inflammatory marker expression, including IL-6, CRP and TNF-three proteins previously linked to MDD caseness. In the instance of IL-6, 18% of the variance was explained by BMI. The exclusion of BMI in previous analyses may have led to false positive associations between IL-6 and MDD case/control status, as MDD cases on average have a higher BMI (particularly in atypical MDD). Indeed, recent meta-analyses revealed that the effect size denoting the association between IL-6 levels and MDD was five times higher when combining results from studies where BMI was not accounted for, suggesting BMI plays a pivotal role in inflating the association between inflammatory markers and MDD.^20^ The relationship between IL-6 and MDD has further been investigated by Mendelian randomisation analyses, showing that genetic variants which alter IL-6 levels are not causally associated with MDD.^49^

Fourth, despite finding that no individual inflammatory marker was powerfully associated with MDD symptomology, our machine learning results suggest inflammatory data may still confer added predictive value in terms of diagnosis and prognosis. We tested the combinatorial influence of our inflammatory marker data using machine learning and found that both within our cross-validation test and our replication tests, inflammatory markers combined with confounders outperformed the predictive capabilities of confounders alone, *Figure 4*. This suggests that information relating to inflammatory markers may improve the precision of diagnoses by 5% when including this information alongside other clinical factors/confounder factors. More striking was that the precision to blindly discriminate an MDD patient in a current episode, increased by 15% when including inflammatory marker data alongside clinical factors (the confounders we examined), *Figure 4*. Unlike IL-16, this finding could be important clinically as it confers a substantially higher precision for discriminating those in a current depressed episode. If these inflammatory measures also precede the onset of a depressed episode, this could be used to initiate rapid treatment before depression symptoms fully present themselves. Thus, although our results show that clinical information/confounders such as BMI do strongly influence inflammatory marker expression, including inflammatory information still added independent information capable of improving diagnosis and prognosis of MDD. Future studies should focus on refining and replicating our results, particularly those variables with high attributive merit scores, *Figure 5*.

While we acknowledge the strengths of the study, which include strict quality control, outlier removal, the screening of control subjects, the randomisation of cases and controls across batches, controlling for batch and study differences, and considering a number of potential confounding factors on inflammatory marker expression, we should also consider the study’s weaknesses. The main weakness of the study is the fact that controls were sampled from a single cohort (SELCoH) that also contained cases, whereas the three additional studies (HDAO, LNBI and GENDEP) were case-only. For each of these four studies there were slightly different blood collection protocols and serum had been stored at −80°C for different periods of time. The population within each sample differed (US, UK only, Europe), and depression caseness was defined slightly differently across the cohorts (but all met criteria for major depressive disorder). Although we corrected for study effects which were consistently significant across our analyses, we may have over- or under-corrected for these differences. Nevertheless, we did correct for study differences and considered this as an appropriate action which was not overly conservative. Furthermore, a significant proportion of our cases were derived from the same cohort as our controls (SELCoH), allowing us to correct for cohort batch effects effectively.

To conclude, our study suggests that there may be no single inflammatory marker predictors of MDD caseness, current episode or depression severity which would be clinically useful. Strong confounding influences such as BMI may have driven previous associations between cytokines such as IL-6 and MDD. Nevertheless, a machine learning approach incorporating clinical/confounder data alongside inflammatory marker data may have some usefulness in improving MDD discrimination, particularly when an individual is currently in a depressed episode, however this requires further validation.

### Conflict of interest

NH has participated in clinical trials sponsored by Lundbeck, Takeda, GlaxoSmithKline, and Pfizer. DS has served on advisory boards for, and received unrestricted grants from Lundbeck and AstraZeneca. AF and PM have received honoraria for participating in expert panels for Lundbeck and GlaxoSmithKline. HW and DC work for Eli Lilly. GB has received consultancy fees and funding from Eli Lilly.

## Acknowledgements

TRP, RC, AK, CG, SLH, MH, and GB are supported by the National Institute for Health Research (NIHR) Biomedical Research Centre for Mental Health at South London and Maudsley NHS Foundation Trust and [Institute of Psychiatry, Psychology & Neuroscience] King’s College London. TRP is also funded by a Medical Research Council Skills Development Fellowship (MR/N014863/1). RU is supported by the Canada Research Chairs Program (file number 950-225925) and the Canadian Institutes of Health Research (funding reference numbers 124976, 142738 and 148394).

The current study was funded by an Eli Lilly LRAP grant awarded to GB, RU, HW, DC and TRP. The Genome-Based Therapeutic Drugs for Depression (GENDEP) study was funded by a European Commission Framework 6 grant (EC Contract Ref LSHB-CT-2003-503428). Lundbeck provided both nortriptyline and escitalopram free of charge for the GENDEP study and GlaxoSmithKline contributed to add-on projects at the London centre. This paper represents independent research funded by the National Institute for Health Research (NIHR) Biomedical Research Centre at South London and Maudsley NHS Foundation Trust and King’s College London. Phase 3 of the SELCoH study was also funded by the Maudsley Charity. The views expressed are those of the authors and not necessarily those of the NHS, the NIHR or the Department of Health. The funders had no role in the design and conduct of the study, in data collection, analysis, or interpretation, or in writing the report.

We would like to acknowledge the work of Prof Peter McGuffin and Anne Farmer for their leading contribution to the GENDEP project.

## Supplementary information

S1: A correlation matrix describing the relationship between inflammatory markers.

S2: Regression results from our case-control comparisons.

S3: Regression results from cases versus ‘super controls’.

S4: ReliefF results describing the contribution of each variable to our machine learning results.

## References

1. Van Deuren M, Dofferhoff ASM, Van Der Meer JWM. Cytokines and the response to infection. J Pathol 2005; 168: 349–356.

2. Imanishi J. Expression of cytokines in bacterial and viral infections and their biochemical aspects. J Biochem 2000; 127: 525–530.

3. Cooper AM, Mayer-Barber JD, Sher A. Role of innate cytokines in mycobacterial infection. Mucosal Immunol 2011; 4: 252–260.

4. Akira S, Kishimoto T. IL-6 and NF-IL6 in acute-phase response and viral infection. Immunol Rev 2002; 127: 25–50.

5. Duque GA, Descoteaux A. Macrophage Cytokines: Involvement in Immunity and Infectious Diseases. Front Immunol 2014; 5: 491.

6. Ridker PM, Hennekens CH, Buring JE, Rifai N. C-reactive protein and other markers of inflammation in the prediction of cardiovascular disease in women. N Engl J Med 2000; 342: 836–843.

7. Murray PJ. The primary mechanism of the IL-10-regulated antiinflammatory response is to selectively inhibit transcription. Proc Natl Acad Sci U S A 2005; 102: 8686–8691.

8. Couper KN, Blount DG, Riley EM. IL-10: The Master Regulator of Immunity to Infection Front Immunol 2014; 105: 141–150.

9. Esser N, Legrand-Poels S, Piette J, Scheen AJ, Paquot N. Inflammation as a link between obesity, metabolic syndrome and type 2 diabetes. Diabetes Res Clin Pract 2014; 105: 141–150.

10. Coussens LM, Werb Z. Inflammation and cancer. Nature 2002; 420: 860–867.

11. Martin C, Tansey KE, Schalkwyk LC, Powell TR. The inflammatory cytokines: molecular biomarkers for major depressive disorder? Biomark Med 2015; 9: 169–180.

12. Lanquillon S, Krieg JC, Bening-Abu-Shach U, Vedder H. Cytokine production and treatment response in major depressive disorder. Neuropsychopharmacol 2000; 22: 370–379.

13. Clerici M, Arosio B, Mundo E, Cattaneo E, Pozzoli S, Dell’osso B, et al. Cytokine polymorphisms in the pathophysiology of mood disorders. CNS Spectr 2009; 14: 419–425.

14. Wium-Andersen MK, Ørsted DD, Nielsen SF, Nordestgaard BG. Elevated C-Reactive Protein Levels, Psychological Distress, and Depression in 73 131 Individuals. JAMA Psychiatry 2013; 70: 176–184.

15. Danner M, Kasl SV, Abramson JL, Vaccarino V. Association between depression and elevated C-reactive protein. Psychosom Med 2003; 65: 347–356.

16. Maes M, Meltzer HY, Bosmans E, et al. Increased plasma concentrations of interleukin-6, soluble interleukin-6, soluble interleukin-2 and transferrin receptor in major depression. J Affect Disord 1995; 34: 301–309.

17. Maes M, Bosmans E, Jongh RD, Kenis G, Vandoolaeghe E, Neels H. Increased serum IL-6 and IL-1 receptor antagonist concentrations in major depression and treatment resistant depression. Cytokine 1997; 9: 853–858.

18. O’Brien SM, Scully P, Fitzgerald P, Scott LV, Dinan TG. Plasma cytokine profiles in depressed patients who fail to respond to selective serotonin reuptake inhibitor therapy. J Psychiatr Res 2007; 41: 326–331.

19. Dowlati Y, Herrmann N, Swardfager W, Liu H, Sham L, Reim EK, et al. A meta-analysis of cytokines in major depression. Biol Psychiatry 2010; 67: 446–457.

20. Howren MB, Lamkin DM, Suls J. Associations of depression with C-reactive protein, IL-1, and IL-6: a meta-analysis. Psychosom Med 2009; 71: 171–186.

21. Opel N, Redlich R, Grotegerda D, Dohm K, Heindel W, Kugel H, et al. Obesity and major depression: Body-mass index (BMI) is associated with a severe course of disease and specific neurostructural alterations. Psychoneuroendocrinology 2015; 51: 219–226.

22. Munafò MR, Araya R. Cigarette smoking and depression: a question of causation. Br J Psychiatry 2010 196: 425–426.

23. Dunlop DD, Song J, Lyons JS, Manheim LM, Chang R. Racial/Ethnic Differences in Rates of Depression Among Preretirement Adults. Am J Public Health 2003 93: 1945–1952.

24. Riolo SA, Nguyen TA, Greden JF, King CA. Prevalence of Depression by Race/Ethnicity: Findings From the National Health and Nutrition Examination Survey III. Am J Public Health 2005; 95: 998–1000.

25. Larsson A, Carlsson L, Lind AL, Gordh T, Bodolea C, Kamali-Moghaddam M, et al. The body mass index (BMI) is significantly correlated with levels of cytokines and chemokines in cerebrospinal fluid. Cytokine 2015; 76: 514–518.

26. Tappia PS, Troughton KL, Langley-Evans SC, Grimble RF. Cigarette smoking influences cytokine production and antioxidant defences. Clin Sci (Lond) 1995; 88: 485–489.

27. Stowe RP, Peek MK, Cutchin MP, Goodwin JS. Plasma Cytokine Levels in a Population-Based Study: Relation to Age and Ethnicity. J Gerontol A Biol Sci Med Sci 2010; 65A: 429–433.

28. Powell TR, Schalkwyk LC, Heffernan AL, et al. Tumor Necrosis Factor and its targets in the inflammatory cytokine pathway are identified as putative transcriptomic biomarkers for escitalopram response. Eur Neuropsychopharmacol 2012; 23: 1105–1114.

29. Powell TR, Tansey KE, Breen G, et al. ATP-binding cassette sub-family F member 1 (ABCF1) is identified as a putative therapeutic target of escitalopram in the inflammatory cytokine pathway. J Psychopharmacol 2013; 27: 609–615.

30. Hatch SL, Frissa S, Verdecchia M, et al. 35. Identifying socio-demographic and socioeconomic determinants of health inequalities in a diverse London community: the South East London Community Health (SELCoH) study. BMC Public Health 2011; 11: 861.

31. Hatch SL, Gazard B, Williams DR, Frissa S, Goodwin L, SELCoH Study Team, et al. Discrimination and common mental disorder among migrant and ethnic groups: findings from a South East London Community sample. Soc Psychiatry Psychiatr Epidemiol 2016; 1: 689–701.

32. Brown J, Evans-Lacko S, Aschan L, et al. Seeking informal and formal help for mental health problems in the community: a secondary analysis from a psychiatric morbidity survey in South London. BMC Psychiatry 2014; 14: 275.

33. Morgan C, Reininghaus U, Reichenberg A. Adversity, cannabis use and psychotic experiences: evidence of cumulative and synergistic effects. Br J Psychiatry 2014; 204: 346–353.

34. Lewis G, Pelosi AJ, Araya R, Dunn G. Measuring psychiatric disorder in the community: a standardized assessment for use by lay interviewers. Psychological Medicine 1992; 22: 465–486.

35. Sheldon RC, Williamson DJ, Corya SA. Olanzapine/fluoxetine combination for treatment-resistant depression: a controlled study of SSRI and nortriptyline resistance. J Clin Psychiatry 2005; 66: 1289–1297.

36. American Psychiatric Association. Diagnostic and statistical manual of mental disorders, fourth edition, text Revision (DSM-IV-TR). 2000. Washington, DC: American Psychiatric Association.

37. Ball SG, Atkinson S, Sparks J, Bangs M, Goldbergr C, Dubé S. Long-Term, Open-Label, Safety Study of Edivoxetine 12 to 18 mg Once Daily as Adjunctive Treatment for Patients With Major Depressive Disorder Who Are Partial Responders to Selective Serotonin Reuptake Inhibitor Treatment. J Clin Psychopharmacol 2015; 35: 266–272.

38. Busner J, Targum SD. The Clinical Global Impressions Scale: Applying a Research Tool in Clinical Practice. Psychiatry (Edgmont) 2007; 4: 28–37.

39. Montgomery, SA, Asberg M. A new depression scale designed to be sensitive to change. Br J Psychiatry 1979; 134: 382–389.

40. Uher R, Farmer A, Maier W, Rietschel M, Hauser J, Marusic A, et al. Measuring depression: comparison and integration of three scales in the GENDEP study. Psychol Med 2008; 38: 289–300.

41. Uher R, Huezo-Diaz P, Perroud N, Smith R, Rietschel M, Mors O, et al. Genetic predictors of response to antidepressants in the GENDEP project. Pharmacogenomics J 2009; 9: 225–233.

42. Uher R, Perroud N, Ng MYM, Hauser J, Henigsberg N, Maier W, et al. Genome-Wide pharmacogenetics of antidepressant response in the GENDEP project. Am J Psychiatry 2010; 167: 555–564.

43. Uher R, Tansey KE, Dew T, Maier W, Mors O, Hauser J, et al. An inflammatory biomarker as a differential predictor of outcome of depression treatment with escitalopram and nortriptyline. Am J Psychiatry. 171: 1278–1286.

44. Hodgson K, Tansey KE, Powell TR, Coppola G, Uher R, Zvezdana Dernovšek M, et al. Transcriptomics and the mechanisms of antidepressant efficacy. Eur Neuropsychopharmacol. 2016; 26: 105–112.

45. World Health Organisation. International Classification of Diseases, 10th Revision (ICD-10). 1992. Geneva: World Health Organisation.

46. Benjamini Y, Hochberg Y. Controlling for the false discovery rate: A practical and powerful approach to multiple testing. JR Stat Soc Series B Stat Methodol 1995; 57: 289–300.

47. Kononenko, I. Estimating Attributes: Analysis and Extensions of RELIEF. European Conference on Machine Learning. 1994; 171–182.

48. Powell TR, Fernandes C, Schalkwyk LC. Depression-Related Behavioral Tests. Curr Protoc Mouse Biol 2012; DOI:10.1002/9780470942390.mo110176.

49. Hyde CL, Nagle MW, Tian C, Chen X, Paciga SA, Wendland JR, et al. Identification of 15 genetic loci associated with risk of major depression in individuals of European descent. Nat Genetics 2016; 48: 1031–1036.

50. Cassano P, Bui E, Rogers AH, Walton ZE, Ross R, Zeng M, et al. Inflammatory cytokines in major depressive disorder: A case-control study. Aust N Z J Psychiatry 2016; 51: 23–31.

